# Neonatal exposure to SARS-CoV-2 spike protein in mice is associated with localized changes in cerebral cortex cell proliferation

**DOI:** 10.64898/2026.11.13.766463

**Authors:** Débora M. Portela, Bruna Andrade, Emanuelle V. De Lima, Raissa R. Christoff, Carolina Batista, Sanclayver Araujo, Leda Castilho, Renato S. Carvalho, Claudia P. Figueiredo, Patrícia P. Garcez, Julia R. Clarke

## Abstract

Coronavirus disease 2019 (COVID-19) can cause acute and persistent neurological manifestations, and evidence suggests these effects may be at least partly explained by direct effects of Severe Acute Respiratory Syndrome Coronavirus 2 (SARS-CoV-2) spike protein, which includes neuroinflammation, cognitive dysfunction, and persistent pain. Because the developing brain undergoes finely regulated processes of cellular proliferation, differentiation, and cerebrovascular maturation, exposure to SARS-CoV-2 or its components during this critical period may interfere with neurodevelopment. We therefore investigated whether neonatal exposure to the SARS-CoV-2 spike protein affects processes underlying brain development. To this end, we used a mouse model previously developed by our group, where Swiss mice received an injection (s.c.) of spike protein from the ancestral viral strain or vehicle at postnatal day 1 (P1). Body weight, body length, and the righting reflex were evaluated until postnatal day 20 (P20). At P10, mouse brains were analyzed for cell proliferation, neural cell populations, and cerebrovascular organization, while blood–brain barrier (BBB) permeability was assessed at P20 using Evans Blue. Neonatal spike protein exposure did not affect body weight, body length, or postural righting reflex. At P10, cell proliferation in spike-exposed animals was comparable to control groups, whereas region-specific alterations in SOX2-positive cells and TBR1-positive neurons were observed. Spike protein exposure did not broadly alter cerebrovascular architecture or BBB permeability, although increased IB4 fluorescence was detected in the lateral dentate gyrus (DG). Our findings provide evidence that exposure to the SARS-CoV-2 spike protein during postnatal development can influence specific aspects of neural development, highlighting the importance of understanding how viral components may affect the developing brain during critical windows of vulnerability.

## 1. Introduction

Coronavirus disease 2019 (COVID-19) has been associated with a broad spectrum of neurological manifestations, ranging from acute symptoms such as anosmia and headache to cerebrovascular and thrombotic complications, as well as persistent symptoms including cognitive dysfunction (“brain fog”) and chronic pain (Bauer et al., 2022; Blomberg et al., 2021; Chou et al., 2021; Davis et al., 2023; Douaud et al., 2022; Greenhalgh et al., 2024; Oba et al., 2026). Increasing evidence suggests that, in addition to its canonical role in viral entry, the Severe Acute Respiratory Syndrome Coronavirus 2 (SARS-CoV-2) spike protein may contribute to neurological and neuroinflammatory alterations (de Lima Almeida et al., 2026; Fontes-Dantas et al., 2023; Jiang et al., 2024; Portela et al., 2026; Schreiber et al., 2025). The spike protein mediates viral entry through binding to angiotensin-converting enzyme 2 (ACE2) and interactions with host proteases, including transmembrane serine protease 2 (TMPRSS2) (Yang & Rao, 2021). Furthermore, the S1 subunit of the spike protein has been shown to cross the blood–brain barrier (BBB) and accumulate in the brain of mice after systemic administration (Rhea et al., 2020). Independently of productive viral infection, extracellular spike protein can also engage innate immune pathways.

This protein acts as a pathogen-associated molecular pattern (PAMP) that can activate Toll-like receptor 4 (TLR4), thereby promoting neuroinflammatory responses (Fontes-Dantas et al., 2023; Frank et al., 2021; Olajide et al., 2021; Zhao et al., 2021). Spike protein-induced TLR4 signaling has been associated with increased microglial reactivity, synaptic alterations, and late cognitive decline, providing a potential mechanism linking exposure to the spike protein with neurological manifestations during and after exposure (Fontes-Dantas et al., 2023). TLR4-dependent signaling has also been implicated in mechanisms underlying persistent pain following exposure to the spike protein (de Lima Almeida et al., 2026).

These effects may be particularly relevant during pregnancy, when the developing nervous system undergoes tightly regulated processes of cell proliferation, differentiation, migration, synaptogenesis, and vascular maturation (Zeiss, 2021). Maternal infection can expose the developing fetus to a systemic inflammatory environment capable of altering neurodevelopment, even in the absence of direct viral infection of the fetal brain (Elgueta et al., 2022; Estes & McAllister, 2016). Consistent with this possibility, maternal SARS-CoV-2 infection has been associated with alterations in glial homeostasis (McMahon et al., 2023) and cerebrovascular integrity in the developing brain (Massimo et al., 2023), and epidemiological studies have raised concerns regarding potential long-term neurological consequences in offspring following maternal infection (Griffin et al., 2026; Martinez et al., 2024; Weiner et al., 2026).

Vertical transmission of SARS-CoV-2 is considered uncommon, with an estimated rate of approximately 7% among infected pregnant individuals (Male, 2022). Notably, the spike protein has been detected on the fetal side of the placenta following maternal SARS-CoV-2 infection (Parcial et al., 2022; Wardhana et al., 2024), raising the possibility that the developing fetus may be exposed to viral components without undergoing productive viral infection. Together, these findings raise the possibility that fetal exposure to SARS-CoV-2 spike protein may represent one mechanism through which maternal infection could influence the developing nervous system.

We previously showed that isolated exposure to SARS-CoV-2 spike protein during the neonatal period induces acute and persistent neuroinflammatory alterations, including increased glial reactivity and changes in synaptic markers (Portela et al., 2026). Here, we assessed the effects of neonatal exposure to the full-length spike protein from the SARS-CoV-2 ancestral strain on early postnatal growth, neural cell proliferation and differentiation, cerebrovascular organization, and BBB permeability. By characterizing these processes during early postnatal development, we aimed to determine whether exposure to the spike protein is sufficient to induce structural or cellular alterations in the developing brain in the absence of productive viral infection.

## 2. Materials and Methods

### Animals and spike infusion

All procedures involving animals were conducted in accordance with the ARRIVE guidelines and applicable institutional and national regulations. The study was approved by the Institutional Animal Care and Use Committee of the Federal University of Rio de Janeiro (UFRJ), Brazil (protocol no. 083/21). Investigators involved in experimental procedures and data analysis were blind to treatment allocation. The recombinant ectodomain of the spike protein derived from the original SARS-CoV-2 Wuhan strain (amino acids 1–1208; 180 kDa) was produced in HEK293 cells and purified in its trimeric prefusion conformation by the Cell Culture Engineering Laboratory (LECC), COPPE/UFRJ, Brazil, as previously described (Fontes-Dantas et al., 2023; Portela et al., 2026; Wrapp et al., 2020). Naïve female Swiss mice aged 10 weeks were obtained from the institutional animal facility and housed for mating for 2 weeks at a ratio of four females per male. Following mating, pregnant females were individually housed until parturition. On postnatal day 1 (P1), litters were standardized to a maximum of 10 pups per dam, whenever possible, with equal numbers of male and female offspring. On P1, pups received a single subcutaneous (s.c.) injection of 10 µL of recombinant SARS-CoV-2 spike protein (1 µg/µL; spike group) or an equivalent volume of phosphate-buffered saline (PBS; vehicle (Veh) group). To minimize potential cross-litter effects, all pups within a given litter received the same treatment. Survival was monitored daily. Body weight was recorded at P3, P10, and P20, and body length (mm) was measured at P10 as the distance from the tip of the nose to the base of the tail.

### Dot Blot Analysis

The presence of spike protein in brain tissue was evaluated by dot blot analysis at multiple time points following exposure. Brain samples were collected from mice in the spike and Veh groups at 1 h, 3 h, 24 h, and 10 days after subcutaneous (s.c.) administration. Tissue samples were homogenized in the presence of protease and phosphatase inhibitors and appropriately diluted before direct application onto nitrocellulose membranes. Membranes were incubated with a polyclonal anti-Spike primary antibody (pAb; Rhea Biotech, 1 mg/mL, 1M-0828, lot no. 20071; 1:4,000) for 1 h at room temperature. Following washing to remove unbound primary antibody, membranes were incubated with an anti-rabbit secondary antibody (1:10,000) for 1 h at room temperature. After removing excess secondary antibodies, the membrane was exposed for 5 min to the detection substrate (SuperSignal West Femto Maximum Sensitivity Substrate, Invitrogen) for signal visualization.

### Postural righting reflex test

The postural righting reflex was assessed daily from P3 to P10. Before testing, the home cages containing the dams and pups were transferred to the testing room and allowed to acclimate for 1 h. Individual pups were gently placed in the center of a clean sheet of paper in a prone position, with all four paws in contact with the testing surface. The pup was then manually rotated onto its back, and timing was initiated immediately after release. The latency to regain the prone position, with all four paws contacting the surface, was recorded as time to turn (s). Following each trial, the pup was returned to its home cage, and the paper surface was replaced before testing the next animal.

### Tissue preparation

Tissues were collected at predefined experimental time points according to the subsequent analyses. For immunofluorescence (IF), animals were anesthetized with ketamine (100 mg/kg) and xylazine (8 mg/kg). Following deep anesthesia, intracardiac perfusion was performed first with 0.1 M PBS, followed by 4% paraformaldehyde (PFA). Brains were subsequently removed and post-fixed in 4% PFA for 48 h before being transferred to 0.1 M PBS for storage. For IF analyses, brains were embedded in 4% agarose and sectioned coronally at 90 μm using a vibratome (Leica, Germany). Sections were maintained free-floating in 0.1 M PBS supplemented with 0.1% sodium azide until immunostaining. For analysis by dot blot, animals were euthanized by cervical dislocation, and brains were rapidly collected and stored at ™80 °C until further processing. For Evans blue assessment of vascular permeability, animals underwent intracardiac perfusion with 0.1 M PBS, after which the brains were collected for subsequent analysis.

### Immunofluorescence assay

IF was used to assess cellular and tissue morphology, quantify specific cell populations, and evaluate the distribution of selected proteins. Brain samples were collected at P10 from animals that had received spike protein or vehicle at P1. Coronal sections (90 μm) were prepared using a vibratome (Leica, Germany). For free-floating immunostaining, sections were subjected to heat-mediated antigen retrieval in citrate buffer (pH 6.0) at 96 °C for 30 min. Sections were subsequently rinsed in 0.1 M PBS containing 0.3% Triton X-100 and blocked for 1 h with 10% normal goat serum (NGS) to minimize nonspecific antibody binding. Tissue sections were then incubated with the primary antibody overnight at 4 °C under gentle agitation (1:500 PH3-rabbit, ref. 06-570; 1:200 SOX2-rabbit, ref. AB5603; 1:200 TBR1-rabbit, ref. AB31940; 1:400 Isolectin B4 (IB4)-488, ref. I214111). Details of all primary and secondary antibodies are provided in Table 1. After primary antibody incubation, sections were extensively washed and incubated for 2 h with the appropriate fluorophore-conjugated secondary (1:500 Alexa 488 goat anti-rabbit, ref. AP132JA4; 1:500 Alexa 546 goat anti-rabbit, ref. A11035). Nuclei were counterstained for 15 min using DAPI nucleic acid stains (Invitrogen). Following additional washing, sections were mounted with Fluoromount mounting medium (Invitrogen) and then covered with a glass coverslip. Images were acquired using a TCS-SPE confocal microscope (Leica, Germany). Cell numbers were determined with the Cell Counter plugin in FIJI/ImageJ, and fluorescence intensity measurements were performed using FIJI/ImageJ. Blood vessel parameters were quantified with AngioTool (Zudaire et al., 2011). For each animal, three brain sections were analyzed, and measurements obtained from the three sections were averaged to generate a single biological replicate. Cell numbers counted were normalized to the mean value of the control group and expressed as a percentage of control.

**Table 1.** Antibodies used for IF.

| Antibody | Source | Dilution | Identifier |
|---|---|---|---|
| **Primary antibodies** |  |  |  |
| pAb Anti-Spike | Rhea Biotech | (1:4,000) | Ref. 1M-0828 |
| Rabbit Anti-PH3 | Sigma-Aldrich | (1:500) | Ref. 06-570 |
| Rabbit Anti-SOX2 | Sigma-Aldrich | (1:200) | Ref. AB5603 |
| Rabbit Anti-TBR1 | Abcam | (1:200) | Ref. AB31940 |
| Isolectin GS-IB4, Alexa Fluor 488 Conjugate | Invitrogen | (1:400) | Ref. I21411 |
| **Secondary antibodies** |  |  |  |
| Horseradish Peroxidase-Conjugated Anti-Rabbit | Abcam | (1:10,000) | Ref. 6721 |
| Alexa 488 goat anti-rabbit | Millipore | (1:500) | Ref. AP132JA4 |
| Alexa 546 goat anti-rabbit | Thermo-Fischer | (1:500) | Ref. A11035 |

### Blood–Brain Barrier Integrity Assessment

Blood–brain barrier (BBB) integrity was assessed at P20 using Evans blue extravasation. Mice that had been exposed to spike or vehicle during the neonatal period were administered 100 µL of 2% Evans blue intravenously via the tail vein. One hour after injection, the animals were intracardiac perfused with 0.1 M PBS, and the brains were collected, weighed, and individually transferred to 2 mL microcentrifuge tubes containing 500 µL formamide. Samples were incubated at room temperature for 48 h to allow extraction of Evans blue from the tissue. The tissue was subsequently removed, and the remaining solution was transferred to a 96-well plate. Evans blue content was quantified spectrophotometrically by measuring absorbance at 620 nm.

### Data analysis

Statistical analyses were conducted using GraphPad Prism 8 (GraphPad Software, CA, USA). Data are presented as mean ± SEM. Potential outliers were screened using the ROUT method with a false discovery rate threshold of Q = 1%, and observations identified as outliers were excluded before subsequent analyses. The distribution of the data was evaluated using the Shapiro–Wilk normality test. For comparisons between two groups, unpaired Student’s *t* tests were used when variances were comparable, whereas the Mann–Whitney U test was applied when the assumptions of equal variance were not met. For experiments involving two independent factors, data were analyzed by two-way analysis of variance (ANOVA) with Sidak’s multiple comparisons test. Statistical significance was defined as p < 0.05.

## 3. Results

SARS-CoV-2 infection during pregnancy has been associated with neuroinflammatory alterations in the offspring (McMahon et al., 2023). We previously showed that exposure to the SARS-CoV-2 spike protein during the neonatal period can recapitulate aspects of this phenotype, resulting in acute and persistent neuroinflammation (Portela et al., 2026). To investigate whether neonatal exposure to spike protein affects early neurodevelopmental processes, including cell proliferation, differentiation, and vascular development, Swiss mice were subcutaneously injected at P1 with full-length spike protein from the ancestral SARS-CoV-2 strain (1 μg/μL; 10 μL) or vehicle (Fig. 1A).

**Figure 1.**
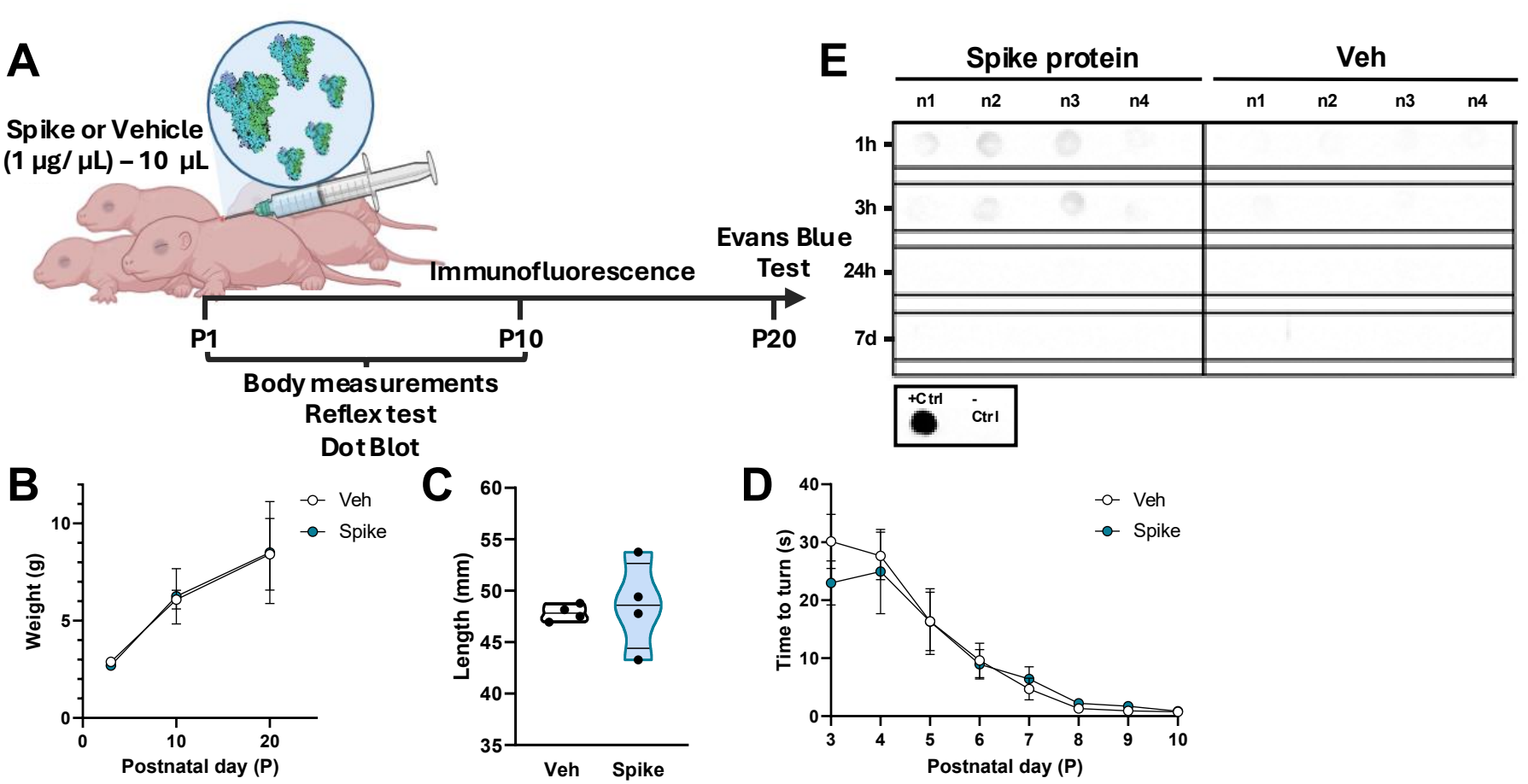
Neonatal exposure to spike protein results in brain uptake but does not alter early postnatal growth or reflex development. (A) Schematic representation of experimental design. Swiss pups received a subcutaneous (s.c.) injection of full-length spike protein (1 µg/µL; 10 µL) or vehicle on postnatal day 1 (P1) and were followed longitudinally for assessment of body weight and length measurements, reflex responses, and spike protein detection by dot blot. Brain samples were collected at P10 for IF analysis and at P20 for assessment of vascular permeability using the Evans blue assay. (B) Body weight measurement at P3, P10, and P20 in mice injected with spike protein or vehicle (n = 4 litters/group; two-way ANOVA). (C) Violin plot showing body length (mm) at P10 in mice injected with spike protein or vehicle (n = 4 litters/group; Student’s *t* test). (D) Time to turn (s) in the righting reflex response test in mice injected with spike protein or vehicle, assessed daily from P3 to P10 (n = 8 per group; two-way ANOVA). (E) Representative dot blots of total brain homogenates from mice exposed to spike protein or vehicle on P1, collected 1 h, 3 h, 24 h, or 7 days after injection. +Ctrl, spike protein; ™Ctrl, buffer used for sample preparation. Data are presented as mean ± S.E.M.

Neonatal spike protein exposure did not affect postnatal body weight gain, assessed at P3, P10, and P20 (Fig. 1B), or body length at P10 (Fig. 1C). Similarly, no differences were observed in the postural righting reflex, assessed daily from P3 to P10 by measuring the time required for pups to return from the supine to the prone position (Fig. 1D). These findings indicate that neonatal spike protein exposure did not produce detectable alterations in early postnatal growth or gross motor development.

To determine the persistence of spike protein in the brain following neonatal exposure, spike protein was assessed by dot blot in whole-brain homogenates at 1 h, 3 h, 24 h, and 7 days after injection (Fig. 1E). Spike protein was detected at 1 and 3 h after exposure but was no longer detectable at 24 h or 7 days. Together, these findings indicate that neonatal spike protein exposure does not alter gross postnatal growth or motor development despite transient detection of spike protein in the brain.

SARS-CoV-2 infection during pregnancy has been associated with alterations in brain structure in the offspring (Weiner et al., 2026). We therefore investigated whether neonatal exposure to the spike protein induces early alterations in brain cellular processes during a critical period of postnatal development. At P10, IF was used to assess cell proliferation, progenitor cells, and neuronal differentiation.

We first assessed cell proliferation by quantifying phospho-histone H3 (PH3)-positive cells. Neonatal spike protein exposure did not significantly affect cell proliferation in the hippocampus, as the number of PH3-positive cells was comparable between spike protein- and vehicle-exposed animals in both the medial dentate gyrus (DG) (Fig. 2A-B) and lateral DG (Fig. 2C-D). Similarly, no significant difference was observed in the number of PH3-positive cells across the whole coronal brain section (Fig. 2E).

**Figure 2.**
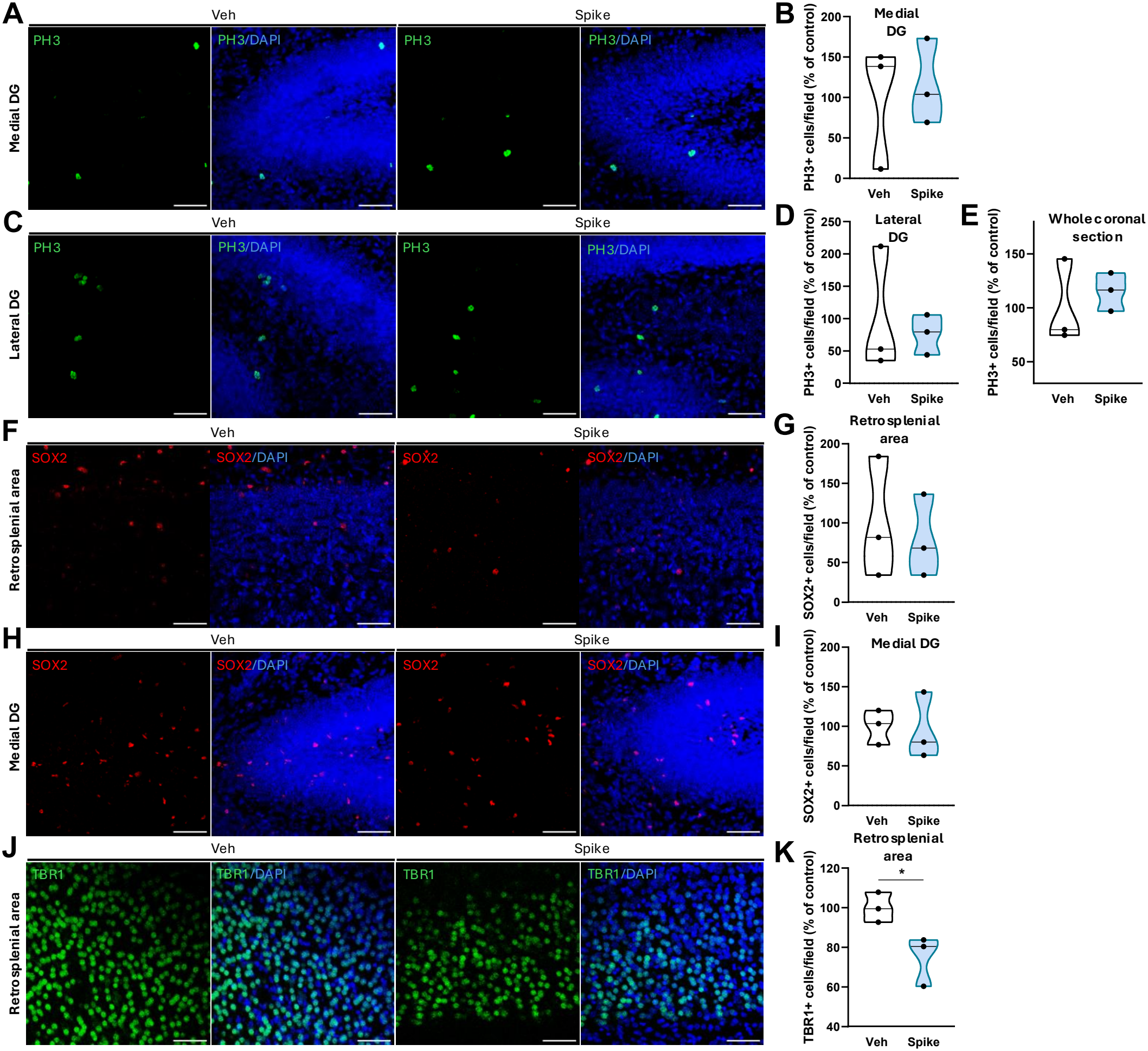
Neonatal exposure to spike protein does not alter cell proliferation or progenitor cell abundance in the hippocampus or cortex but reduces mature neurons in the retrosplenial area at P10. (A) Representative images of PH3 immunolabeling and DAPI staining in the medial dentate gyrus (DG) of the hippocampus at P10 in mice injected with spike protein or vehicle at P1 (scale bar: 50 µm). (B) Violin plot showing the number of PH3-positive cells, expressed as a percentage of the control group, in the medial DG at P10 in mice injected with spike protein or vehicle at P1 (n = 3 per group; Student’s *t* test). (C) Representative images of PH3 immunolabeling and DAPI staining in the lateral DG of the hippocampus at P10 in mice injected with spike protein or vehicle at P1 (scale bar: 50 µm). (D, E) Violin plots showing the number of PH3-positive cells, expressed as a percentage of control group, in the lateral DG (n = 3 per group; Student’s *t* test) and throughout the entire coronal brain section (n = 3 per group; Student’s *t* test), respectively, at P10 in mice injected with spike protein or vehicle at P1. (F) Representative images of SOX2 immunolabeling and DAPI staining in the retrosplenial area at P10 in mice injected with spike protein or vehicle at P1 (scale bar: 50 µm). (G) Violin plot showing the number of SOX2-positive cells, expressed as a percentage of the control group, in the retrosplenial area at P10 in mice injected with spike protein or vehicle at P1 (n = 3 per group; Student’s *t* test). (H) Representative images of SOX2 immunolabeling and DAPI staining in the medial DG of the hippocampus at P10 in mice injected with spike protein or vehicle at P1 (scale bar: 50 µm). (I) Violin plot showing the number of SOX2-positive cells, expressed as a percentage of the control group, in the medial DG at P10 in mice injected with spike protein or vehicle at P1 (n = 3 per group; Student’s *t* test). (J) Representative images of TBR1 immunolabeling and DAPI staining in the retrosplenial area at P10 in mice injected with spike protein or vehicle at P1 (scale bar: 50 µm). (K) Violin plot showing the number of TBR1-positive cells, expressed as a percentage of the control group, in the retrosplenial area at P10 in mice injected with spike protein or vehicle at P1 (n = 3 per group; \**p* = 0.0419; Student’s *t* test). Data are presented as mean ± S.E.M.

Studies previously observed increased microglial abundance and reactivity during neurodevelopment following maternal SARS-CoV-2 exposure (McMahon et al., 2023) and after neonatal spike protein exposure (Portela et al., 2026). Because increased microglial activity has been reported to interfere with neural progenitor cell proliferation and neuronal differentiation (Appel et al., 2018), we next investigated whether neonatal spike protein exposure altered cell populations positive for SOX2, typically used as a marker of neural stem/progenitor cells, at P10. No significant difference in the number of SOX2-positive cells was observed in the retrosplenial cortex (Fig. 2F-G). In contrast, spike protein-exposed animals showed an increase in the number of SOX2-positive cells in the somatosensory cortex (Suppl. Fig. 1A-B). In the hippocampus, the number of SOX2-positive cells did not differ significantly between groups in the medial DG (Fig. 2H-I), lateral DG (Suppl. Fig. 1C-D), or CA1 (Suppl. Fig. 1E-F).

We next examined neuronal differentiation by quantifying TBR1-positive cells, using TBR1 as a marker of developing postmitotic neurons. Neonatal spike protein exposure was associated with a significant reduction in the number of TBR1-positive cells in the retrosplenial cortex at P10 (Fig. 2J-K). However, no significant difference was observed in the number of TBR1-positive cells in the somatosensory cortex (Suppl. Fig. 1G-H).

Together, these findings indicate that neonatal spike protein exposure does not broadly alter cell proliferation or SOX2-positive cell populations across the brain at P10, but is associated with region-specific changes in SOX2-positive cells and a reduction in TBR1-positive neuronal populations in the retrosplenial cortex.

COVID-19 has been associated with neurovascular alterations, including disruption of the BBB, fibrinogen extravasation, immune cell infiltration, and increased glial reactivity (Bocci et al., 2021; Matschke et al., 2020; Schurink et al., 2020; Wenzel et al., 2021). Similarly, intrauterine exposure to SARS-CoV-2 has been associated with vascular disruption and immune cell infiltration in the developing brain (Massimo et al., 2023). We therefore investigated whether neonatal exposure to the spike protein induces early alterations in cerebrovascular density, organization, and BBB integrity. At P10, the cerebrovasculature was visualized by IB4 staining, which labels endothelial cells in blood vessels and can also detect microglial cells, and vascular architecture was quantified using AngioTool analysis.

In the retrosplenial cortex, no significant difference in IB4 fluorescence intensity was observed between spike protein- and vehicle-exposed animals at P10 (Fig. 3A-B). Consistent with this finding, AngioTool analysis revealed no significant differences in vascular area (Fig. 3C-D), average vessel length (Fig. 3E), or number of vessel junctions (Fig. 3F), indicating that neonatal spike protein exposure did not detectably alter vascular density or network organization in the retrosplenial cortex. A similar pattern was observed in the somatosensory cortex, where neither IB4 fluorescence intensity (Suppl. Fig. 2A-B) nor vascular area, average vessel length, or number of junctions differed between groups (Suppl. Fig. 2C–F).

**Figure 3.**
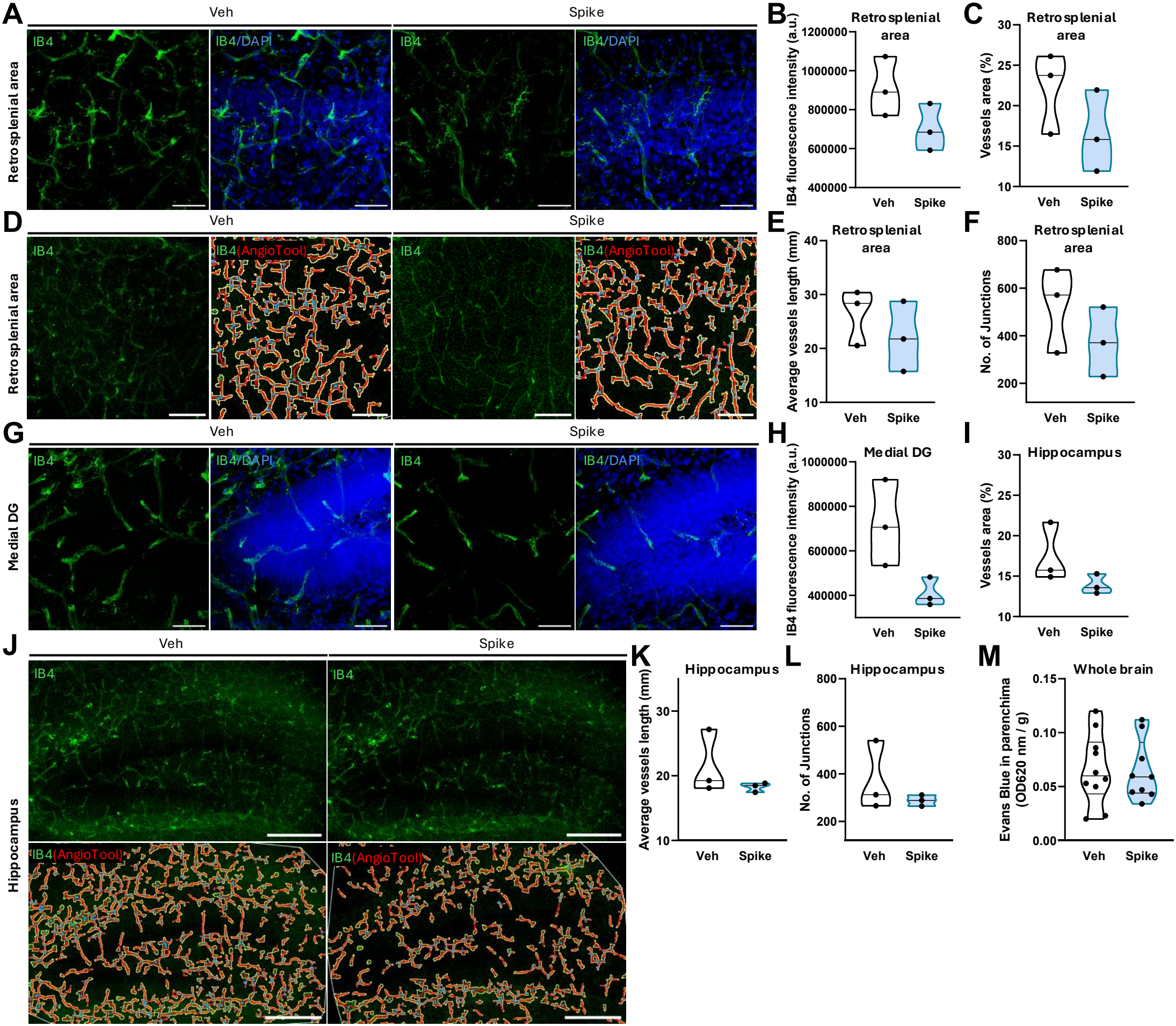
Neonatal exposure to spike protein does not alter vascular parameters in the retrosplenial area or hippocampus at P10. (A) Representative images of IB4 and DAPI staining in the retrosplenial area at P10 in mice injected with spike protein or vehicle at P1 (scale bar: 50 µm). (B) Violin plot showing IB4 fluorescence intensity (arbitrary units, a.u.) in the retrosplenial area at P10 (n = 3 per group; Student’s *t* test). (D) Representative images of IB4 staining with blood vessels traced using AngioTool in the retrosplenial area at P10 (scale bar: 100 µm). (C–F) Violin plots showing vessel area (%; C; n = 3 per group; Student’s *t* test), average vessel length (mm; E; n = 3 per group; Student’s *t* test), and number of junctions (F; n = 3 per group; Student’s *t* test) in the retrosplenial area at P10. (G) Representative images of IB4 and DAPI staining in the medial DG of the hippocampus at P10 in mice injected with spike protein or vehicle at P1 (scale bar: 50 µm). (H) Violin plot showing IB4 fluorescence intensity (a.u.) in the medial DG at P10 (n = 3 per group; Student’s *t* test). (J) Representative images of IB4 staining with blood vessels traced using AngioTool in the hippocampus at P10 (scale bar: 200 µm). (I–L) Violin plots showing vessel area (%; I; n = 3 per group; Student’s *t* test), average vessel length (mm; K; n = 3 per group; Student’s *t* test), and number of junctions (L; n = 3 per group; Student’s *t* test) in the hippocampus at P10. (M) Violin plot showing Evans blue extravasation into the brain parenchyma, assessed 20 days after injection of spike protein or vehicle (n = 10 Veh; n = 9 spike; Student’s *t* test). Data are presented as mean ± S.E.M.

We next assessed the hippocampal vasculature. IB4 fluorescence intensity did not differ between groups in the medial DG (Fig. 3G-H) or CA1 (Suppl. Fig. 2I-J). In contrast, spike protein exposure was associated with a decrease in IB4 fluorescence intensity in the lateral DG (Suppl. Fig. 2G-H). Despite this regional change in IB4 fluorescence, AngioTool analysis revealed no significant differences in vascular area (Fig. 3I-J), average vessel length (Fig. 3K), or number of vessel junctions (Fig. 3L) in the hippocampus at P10.

Finally, to determine whether neonatal spike protein exposure affected BBB integrity at a later developmental stage, we assessed Evans Blue extravasation in the brain at P20. No significant difference in Evans Blue accumulation in the brain parenchyma was observed between spike protein- and vehicle-exposed animals (Fig. 3M), indicating no detectable increase in BBB permeability under these experimental conditions.

Together, these findings indicate that neonatal spike protein exposure does not produce broad alterations in cerebrovascular architecture or BBB permeability during early postnatal development, although a region-specific decrease in IB4 fluorescence was detected in the lateral DG at P10.

## 4. Discussion

COVID-19 is associated with a broad range of neurological manifestations (Bauer et al., 2022; Blomberg et al., 2021; Chou et al., 2021; Davis et al., 2023; Greenhalgh et al., 2024; Oba et al., 2026). Maternal SARS-CoV-2 infection raised particular concern because infections during pregnancy can interfere with brain development through inflammatory and other systemic mechanisms (Kirtsman et al., 2020; Lokken, Huebner, et al., 2021; Lokken, Taylor, et al., 2021; Schoenmakers et al., 2021; Vivanti et al., 2020). Although vertical transmission of SARS-CoV-2 appears to be rare, it has been documented in approximately 7% of pregnancies (Male, 2022). Importantly, in some cases, viral proteins have been detected in placental and fetal tissues even when definitive evidence of vertical transmission was not established (Parcial et al., 2022; Wardhana et al., 2024). These findings raise the possibility that exposure to viral components, even in the absence of productive fetal infection, may influence the developmental environment through inflammation.

To investigate whether exposure to the SARS-CoV-2 spike protein could affect neurodevelopment, we administered 10 μg of full-length spike protein (1 μg/μL) or vehicle subcutaneously to mice at postnatal day 1 (P1). This period corresponds approximately to the late gestational period of human brain development and is characterized by intense cellular proliferation, gliogenesis, synaptogenesis, and cerebrovascular maturation, making the developing brain particularly vulnerable to environmental and inflammatory insults (Workman et al., 2013).

Using this experimental paradigm, we did not observe alterations in general postnatal development, as assessed by body weight gain, body length, and the postural righting reflex. However, neonatal spike exposure was associated with specific neural alterations at P10. Overall cell proliferation was preserved, whereas SOX2-positive progenitor/immature neural cells and TBR1-positive differentiated neurons showed region-specific changes. Specifically, spike protein exposure increased the number of SOX2-positive cells in the somatosensory cortex and decreased the number of TBR1-positive neurons in the retrosplenial area. The absence of changes in PH3-positive proliferating cells at P10, together with the alterations in neural cell populations, may indicate that the effects of neonatal spike exposure are not reflected by a persistent change in proliferation at this time point. Instead, exposure during an earlier developmental window may have transiently influenced progenitor dynamics or subsequent differentiation, ultimately altering the abundance of specific neural populations at P10. Longitudinal analyses of proliferation and differentiation would be necessary to distinguish between these possibilities.

These findings are consistent with growing evidence that shows that SARS-CoV-2 infection and exposure to viral components can affect neural and neuroimmune processes. Previous studies have reported that spike protein exposure can activate inflammatory pathways, alter glial responses, and affect neuronal function, suggesting that the spike protein may contribute to neurological effects beyond those resulting from active viral replication (Frank et al., 2021). In the developing brain, where neuronal differentiation, synaptic maturation, and glial development occur simultaneously, transient inflammatory or cellular disturbances may influence ongoing developmental trajectories (Xu et al., 2024). In agreement with this concept, we previously showed that neonatal exposure to spike protein was associated with a reduction in SATB2-positive neurons in specific cortical regions, accompanied by a pronounced neuroimmune response (Portela et al., 2026). Together, these findings suggest that spike exposure may affect multiple aspects of neural development and that distinct neuronal populations and brain regions may differ in their susceptibility to this stimulus.

Regional differences in cellular composition, developmental timing, vascularization, or sensitivity to inflammatory signaling may contribute to this selective vulnerability (Ahirwar et al., 2025; Bilbo & Schwarz, 2012; Paolicelli et al., 2022; Thion et al., 2018; Zeiss, 2021). Thus, the effects of spike exposure may depend not only on the intensity of the stimulus but also on the developmental state and cellular characteristics of each brain region. The increase in SOX2-positive cells in the somatosensory cortex and reduction in TBR1-positive neurons in the retrosplenial area observed here, together with the previously reported alterations in SATB2-positive neurons, support the possibility that different neuronal populations may respond differently to the same developmental insult.

The neonatal brain is also undergoing extensive angiogenesis and BBB maturation (Ahirwar et al., 2025; Collignon et al., 2024; Vasudevan et al., 2008; Wälchli et al., 2015). Developmental inflammatory insults have been associated with altered vascular maturation, including changes in vessel development (Garcez et al., 2018) and BBB permeability dysfunction (Ahirwar et al., 2025; Wälchli et al., 2015). In contrast, neonatal spike exposure did not produce broad alterations in cerebrovascular organization at P10, and BBB permeability was preserved at P20. Although overall cerebrovascular organization was preserved, neonatal spike exposure was associated with a localized reduction in IB4 fluorescence in the lateral DG. Because IB4 labeling can identify endothelial cells as well as specific populations of microglia, this finding should be interpreted cautiously and does not, by itself, establish a reduction in vascular density or endothelial cell number. Nevertheless, the localized decrease in IB4 fluorescence may indicate a subtle alteration in the cellular or vascular environment of the DG that was not accompanied by detectable changes in broader measures of cerebrovascular architecture. Further characterization using complementary endothelial and vascular markers will be necessary to determine the basis and biological significance of this finding. Together, these results suggest that spike exposure does not cause a broad disruption of cerebrovascular architecture or persistent BBB dysfunction.

Several limitations should be considered when interpreting these findings. Our experimental model used purified full-length spike protein rather than SARS-CoV-2 infection and therefore does not recapitulate the complex interactions associated with viral replication or the systemic immune response to infection. Nevertheless, this approach allowed us to investigate the potential effects of spike protein exposure independently of active viral replication. Importantly, previous studies have shown that the SARS-CoV-2 spike protein, particularly its S1 subunit, can persist in tissues and immune cells for prolonged periods after SARS-CoV-2 infection or spike protein exposure (Patterson et al., 2022; Rhea et al., 2020; Rong et al., 2024). That detection in specific immune cells (Patterson et al., 2022) and the meninges has been associated with neurological manifestations of long COVID (Rong et al., 2024). Although our model did not use the isolated S1 subunit, the full-length spike protein contains the S1 domain. In our experimental model, spike protein was detectable in whole brain tissue at early time points following neonatal administration, including 1 and 3 h after exposure, whereas it was no longer detected at later time points. These findings indicate that the administered protein reaches the brain during the early post-exposure period, but do not provide evidence for its persistence until P10. Finally, because our analyses were performed at specific developmental time points, we cannot determine whether the observed cellular alterations are transient or persistent, nor whether they translate into functional consequences later in life.

Importantly, our findings do not indicate a generalized impairment of postnatal development following neonatal spike exposure. Rather, they suggest that exposure to a viral component during a critical developmental window may produce selective alterations in specific neural populations and brain regions, even in the absence of detectable changes in overall growth, cerebrovascular architecture, or BBB permeability. More broadly, these findings highlight the importance of investigating how exposure to viral components during critical periods of brain maturation may influence specific neurodevelopmental processes and contribute to alterations that may not be apparent from measures of general development alone.

## Author Contributions

Conceptualization: LC, CPF, PPG, JRC; Funding acquisition: CPF, PPG, JRC; Investigation: DMP, BA, EVL, CB, SA, RSC, RRC; Methodology: DMP, CB, SA; Roles/Writing - original draft: DMP, PPG, JRC; Writing - review & editing: all authors.

## Authors declare no conflict of interest

## Declaration of Generative AI and AI-assisted technologies in the writing process

AI-assisted technologies were only used to improve readability and language, and not for text generation or data interpretation.

## Acknowledgements

We thank Ana Claudia Rangel for project management, the ICB Microscopy Unit for access to the Leica SPE Confocal Microscope and the ICB Animal Facility for animal handling and care.

## Figure Legends

**Supplementary Figure 1.**
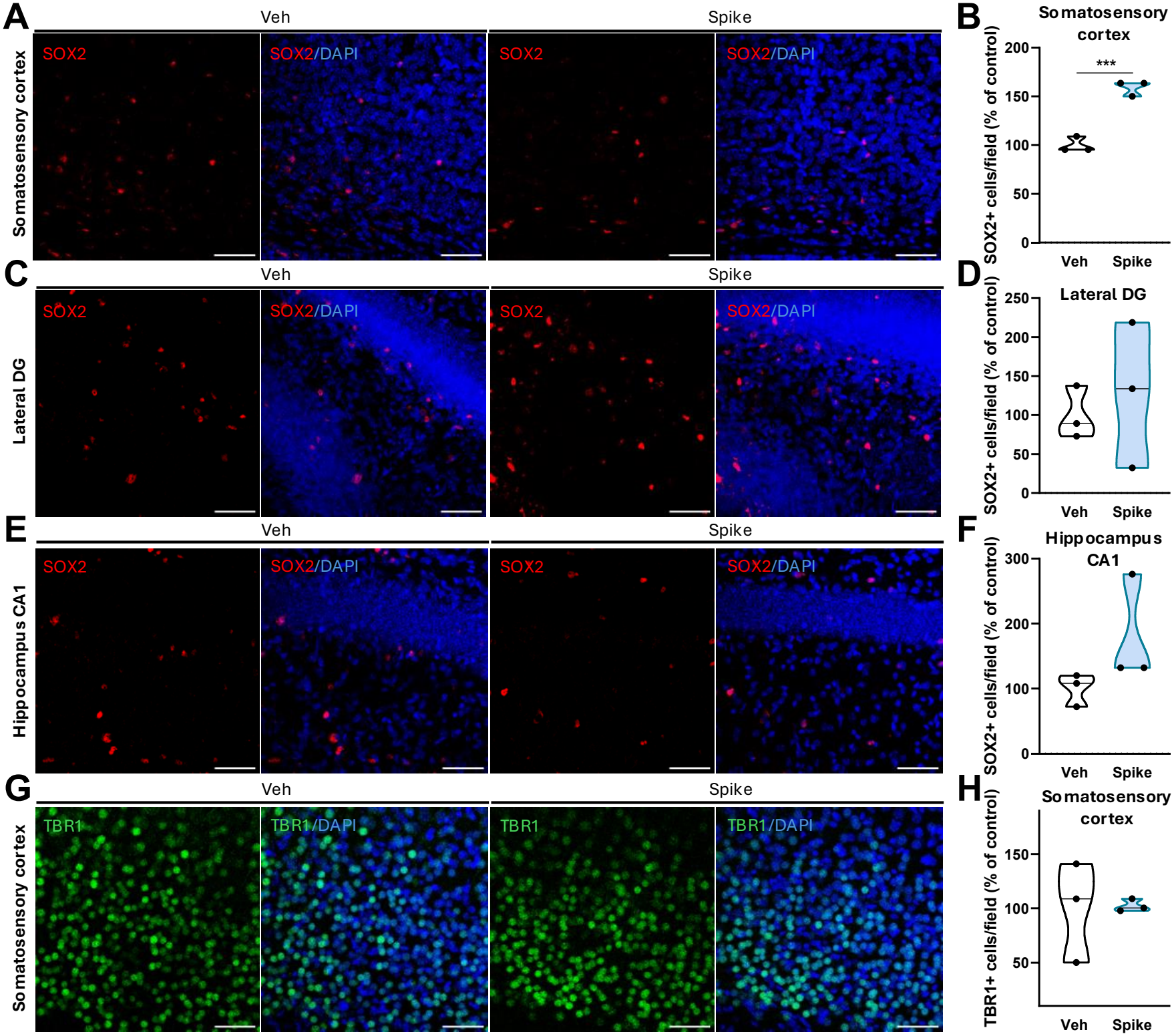
Neonatal exposure to spike protein increases progenitor cell abundance in the somatosensory cortex but does not affect hippocampal progenitor cell or somatosensory-cortical mature neuron abundance at P10. (A) Representative images of SOX2 immunolabeling and DAPI staining in the somatosensory cortex at P10 in mice injected with spike protein or vehicle at P1 (scale bar: 50 µm). (B) Violin plot showing the number of SOX2-positive cells, expressed as a percentage of the control group, in the somatosensory cortex at P10 in mice injected with spike protein or vehicle at P1 (n = 3 per group; \*\*\**p* = 0.0008; Student’s *t* test). (C) Representative images of SOX2 immunolabeling and DAPI staining in the lateral DG of the hippocampus at P10 in mice injected with spike protein or vehicle at P1 (scale bar: 50 µm). (D) Violin plot showing the number of SOX2-positive cells, expressed as a percentage of the control group, in the lateral DG at P10 in mice injected with spike protein or vehicle at P1 (n = 3 per group; Student’s *t* test). (E) Representative images of SOX2 immunolabeling and DAPI staining in the CA1 of the hippocampus at P10 in mice injected with spike protein or vehicle at P1 (scale bar: 50 µm). (F) Violin plot showing the number of SOX2-positive cells, expressed as a percentage of the control group, in the CA1 at P10 in mice injected with spike protein or vehicle at P1 (n = 3 per group; Student’s *t* test). (G) Representative images of TBR1 immunolabeling and DAPI staining in the somatosensory cortex at P10 in mice injected with spike protein or vehicle at P1 (scale bar: 50 µm). (K) Violin plot showing the number of TBR1-positive cells, expressed as a percentage of the control group, in the somatosensory cortex at P10 in mice injected with spike protein or vehicle at P1 (n = 3 per group; Student’s *t* test). Data are presented as mean ± S.E.M.

**Supplementary Figure 2.**
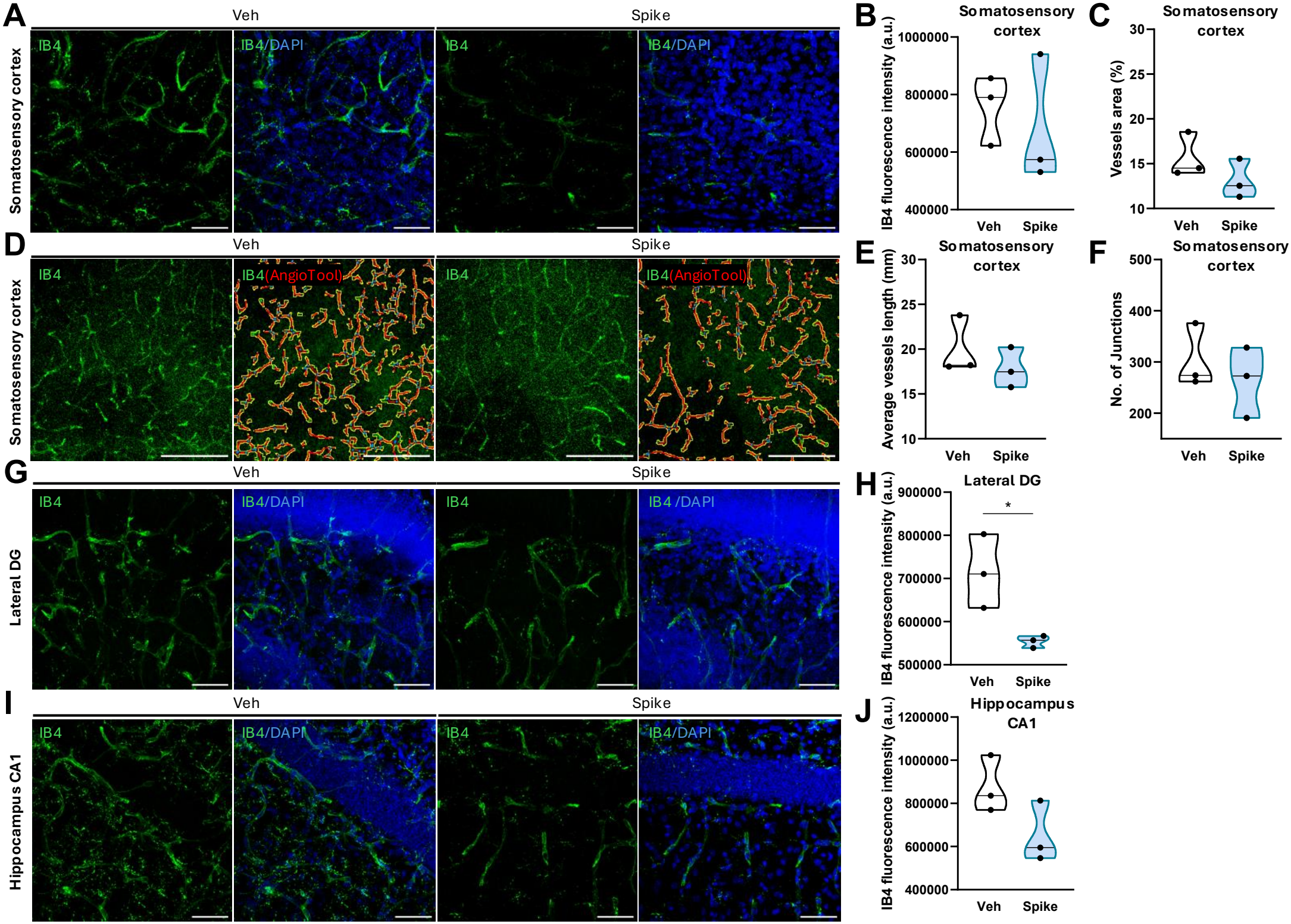
Neonatal exposure to spike protein does not alter vascular parameters in the somatosensory cortex or hippocampus at P10. (A) Representative images of IB4 and DAPI staining in the somatosensory cortex at P10 in mice injected with spike protein or vehicle at P1 (scale bar: 50 µm). (B) Violin plot showing IB4 fluorescence intensity (a.u.) in the somatosensory cortex at P10 (n = 3 per group; Student’s *t* test). (D) Representative images of IB4 staining with blood vessels traced using AngioTool in the somatosensory cortex at P10 (scale bar: 100 µm). (C–F) Violin plots showing vessel area (%; C; n = 3 per group; Student’s *t* test), average vessel length (mm; E; n = 3 per group; Mann-Whitney test), and number of junctions (F; n = 3 per group; Student’s *t* test) in the somatosensory cortex at P10. (G) Representative images of IB4 and DAPI staining in the lateral DG of the hippocampus at P10 in mice injected with spike protein or vehicle at P1 (scale bar: 50 µm). (H) Violin plot showing IB4 fluorescence intensity (a.u.) in the lateral DG at P10 (n = 3 per group; \**p* = 0.0325; Student’s *t* test). (I) Representative images of IB4 and DAPI staining in the CA1 of the hippocampus at P10 in mice injected with spike protein or vehicle at P1 (scale bar: 50 µm). (J) Violin plot showing IB4 fluorescence intensity (a.u.) in the CA1 at P10 (n = 3 per group; Student’s *t* test). Data are presented as mean ± S.E.M.

